## Supplementary Table 1 for "Regulative synthesis of capsular polysaccharides in the pathogenesis of *Streptococcus suis*"

**Supplemental Table 1. Information of the strains used in the study**

| Strain Name | Serotype | Isolated from | | Disease/ Clinical Manifestation | Year of isolation | Reference |
| --- | --- | --- | --- | --- | --- | --- |
|  |  | Host | Sample |  |  |  |
| 05ZYH33 | 2 | Pig | Brain | Diseased | 2005 | 1 |
| 4961 | 3 | Pig |  | Diseased |  | 2 |
| J99 | 7 | Pig | Lung | Pneumonia | 2008 |  |
| D12 | 9 | Pig |  |  |  | 3 |
| SS12 | 1/2 | Pig |  |  |  | 3 |

**Reference**

1. Hu Q, Liu P, Yu Z, et al. Identification of a cell wall-associated subtilisin-like serine protease involved in the pathogenesis of Streptococcus suis serotype 2. Microb Pathog **2010**; 48:103-9.

2. Tien le HT, Nishibori T, Nishitani Y, Nomoto R, Osawa R. Reappraisal of the taxonomy of Streptococcus suis serotypes 20, 22, 26, and 33 based on DNA-DNA homology and sodA and recN phylogenies. Vet Microbiol **2013**; 162:842-9.

3. Zhang A, Yang M, Hu P, et al. Comparative genomic analysis of Streptococcus suis reveals significant genomic diversity among different serotypes. BMC genomics **2011**; 12:523.
